# Validation of genome-wide polygenic risk scores for coronary artery disease in French Canadians

**DOI:** 10.1101/538470

**Authors:** Florian Wünnemann, Ken Sin Lo, Alexandra Langford-Avelar, David Busseuil, Marie-Pierre Dubé, Jean-Claude Tardif, Guillaume Lettre

## Abstract

Coronary artery disease (CAD) represents one of the leading causes of morbidity and mortality worldwide. Given the healthcare risks and societal impacts associated with CAD, their clinical management would benefit from improved prevention and prediction tools. Polygenic risk scores (PRS) based on an individual’s genome sequence are emerging as potentially powerful biomarkers to predict the risk to develop CAD. Two recently derived genome-wide PRS have shown high specificity and sensitivity to identify CAD cases in European-ancestry participants from the UK Biobank. However, validation of the PRS predictive power and transferability in other populations is now required to support their clinical utility. We calculated both PRS (GPS_CAD_ and metaGRS_CAD_) in French-Canadian individuals from three cohorts totaling 3639 prevalent CAD cases and 7382 controls, and tested their power to predict prevalent, incident and recurrent CAD. We also estimated the impact of the founder French-Canadian familial hypercholesterolemia deletion (*LDLR* delta > 15kb deletion) on CAD risk in one of these cohorts and used this estimate to calibrate the impact of the PRS. Our results confirm the ability of both PRS to predict prevalent CAD comparable to the original reports (area under the curve (AUC) = 0.72-0.84). Furthermore, the PRS identified about 6-7% of individuals at CAD risk similar to carriers of the *LDLR* delta > 15kb mutation, consistent with previous estimates. However, the PRS did not perform as well in predicting incident (AUC= 0.56 - 0.60) or recurrent (AUC= 0.56 - 0.60) CAD. This result suggests that additional work is warranted to better understand how ascertainment biases and study design impact PRS for CAD. Collectively, our results confirm that novel, genome-wide PRS are able to predict CAD in French-Canadians; with further improvements, this is likely to pave the way towards more targeted strategies to predict and prevent CAD-related adverse events.

## Introduction

Genome-wide association studies (GWAS) have shed light on the polygenic architecture of human quantitative traits such as height and blood pressure, as well as common diseases such as type 2 diabetes and coronary artery disease (CAD) ^1–4^. These studies have shown that complex human phenotypes are controlled by hundreds of genetic variants, each with small effect size. Although individually they contribute to a small fraction of the phenotypic variation, together they account for a relatively large fraction of the heritability ^5^. This observation has raised the possibility to use genetic variants distributed across the genome to calculate polygenic risk scores (PRS) and use them to predict the risk to develop diseases ^6^. The availability of large human genetic datasets, such as the UK Biobank, now allows for calibration and validation of genome-wide PRS in >100,000 individuals.

CAD remains one of the main causes of morbidity and mortality worldwide ^7^. GWAS have already identified >100 loci associated with CAD, mostly in populations of European ancestry ^2,8^. Early prediction would benefit prevention, optimal management, and treatment strategies for CAD. Although CAD has high heritability (50-60%) ^9,10^, genetic testing is not readily used in the clinic, except in the context of Mendelian disease such as familial hypercholesterolemia (FH). Two recently developed genome-wide PRS for CAD by Khera et *al*. (GPS_CAD_) and Inouye et *al*. (metaGRS_CAD_) suggest that genetic risk prediction for CAD is ready to be applied in the clinical setting ^11,12^. Khera and colleagues utilized the LDpred algorithm to model linkage disequilibrium and variant effect sizes from a CAD GWAS in the UK Biobank to create GPS_CAD_, which includes >6 million genetic variants throughout the genome ^11,13^. In contrast, Inouye and colleagues created a PRS termed metaGRS_CAD_ with >1.7 million variants, themselves explaining 26% of CAD heritability, using a meta-analysis of association results from three large CAD GWAS ^2,14,15^. The conclusions from both studies were encouraging. Khera et *al*. showed that GPS_CAD_ can identify a significant portion of individuals in the general population with a polygenic CAD risk as high as those who carry mutations that cause FH ^11^. For Inouye et *al*., the CAD risk estimated with metaGRS_CAD_ was higher than the risk conferred by any single traditional risk factors such as smoking or hypertension ^12^.

Although these results are promising, the introduction of CAD PRS in clinical practice is likely to encounter resistance^16–18^. In particular, whether PRS are sufficiently accurate to justify on their own early interventions – including pharmaceutical treatments – is an important debate. For this reason, it is critical to validate PRS in additional populations (GPS_CAD_ and metaGRS_CAD_ were only tested in European-ancestry participants from the UK Biobank) and determine whether ascertainment biases and/or study design impact their clinical utility. Here, we validated these two novel CAD PRS in individuals of French-Canadian descent recruited from population- and hospital-based cohorts. We evaluated the performance of these polygenic predictors on prevalent, incident, and recurrent CAD. Finally, we used whole-genome sequence data to identify participants that carry a known FH mutation, and compared its impact on CAD risk with that due to the inheritance of millions of weak effects common variants.

## Results

### Genome-wide PRS for prevalent CAD in French Canadians

Using both models (GPS_CAD_ and metaGRS_CAD_), we calculated PRS in French Canadians from three studies: two hospital-based cohorts from the Montreal Heart Institute (MHI) Biobank (phase 1, n=1,964; phase 2, n=3,309) ^19,20^, and 5,762 participants from CARTaGENE, a public health research platform in the Province of Quebec, Canada ^21^. We present demographics and baseline clinical information for all participants in Table 1. Following DNA genotyping and variant imputation (**Methods**), most variants used to calculate GPS_CAD_ and metaGRS_CAD_ were present in our datasets (missingness range: 0.09-6.96%, **Table S1**), suggesting that our study design can accurately capture the previously proposed CAD polygenic models. Both PRS were strongly correlated with each other in the French-Canadian datasets (Pearson’s *r*>0.73, p-value<2.2×10^-16^) (Figure 1). We tested the association between the CAD PRS and prevalent CAD status in all three cohorts. The distributions of both GPS_CAD_ and metaGRS_CAD_ were shifted towards higher values in CAD cases when compared to controls (Figure 2). Combining results across the three cohorts, we found that one standard deviation increase in GPS_CAD_ or metaGRS_CAD_ was associated with increased odds of CAD of 1.63 (p-value = 7.5×10^-50^) and 1.74 (p-value = 8.5×10^-62^), respectively (Table 2). In terms of prediction of prevalent CAD in French Canadians, the area under the receiving operating characteristic curve (AUC) for both PRS were 0.72-0.84, largely consistent with the original reports (Table 2).

**Table 1.** Demographics and clinical information for the participants involved in the study. Coronary artery disease (CAD) is defined as previous diagnosis of myocardial infarction or revascularization procedures (percutaneous coronary intervention or coronary artery bypass grafting). Hypertension is defined as a previous diagnosis of hypertension, on antihypertensive therapy or with systolic blood pressure ≥140 mmHg or diastolic blood pressure ≥90 mmHg. Diabetes mellitus is defined as a previous diagnosis of diabetes or treatment with antidiabetic drugs. Dyslipidemia is defined as a previous diagnosis of hypercholesterolemia or treatment with lipid-lowering drugs. N.A. = not available.

| Characteristic | MHI Biobank phase 1 |  | MHI Biobank phase 2 |  | CARTaGENE |  |
| --- | --- | --- | --- | --- | --- | --- |
| Genotyping platform | Low-pass WGS (5X) |  | Illumina MEGA |  | Illumina GSA |  |
| Baseline status | *Controls* | *Cases* | *Controls* | *Cases* | *Controls* | *Cases* |
| Sample size, n (% women) | 976 (28) | 974 (27) | 817 (60) | 2492 (17) | 5589 (60) | 173 (17) |
| Mean age, years (standard deviation) | 66.0 (10.1) | 66.9 (8.87) | 66.0 (10.7) | 66.7 (8.31) | 54.9 (7.78) | 60.5 (6.90) |
| Type 2 diabetes, n (%) | 195 (20) | 291 (30) | 59 (7) | 702 (28) | 336 (6) | 42 (24) |
| Hypertension, n (%) | 632 (65) | 751 (77) | 287 (35) | 1878 (75) | 1132 (20) | 94 (54) |
| Dyslipidemia, n (%) | 741 (76) | 923 (95) | 306 (37) | 2350 (94) | 1551 (28) | 121 (70) |
| Mean LDL-cholesterol, mmol/L (standard deviation) | 2.65 (0.87) | 2.09 (0.7) | 3.12 (0.85) | 2.67 (0.8) | 3.03 (0.85) | 1.99 (0.76) |
| Follow-up number of years, median (range) | 4.1 (2.8–6.6) | 4.1 (2.7–7) | 3.8 (0.1–7.2) | 3.7(1.1–7) | N.A. | N.A. |

**Table 2.**
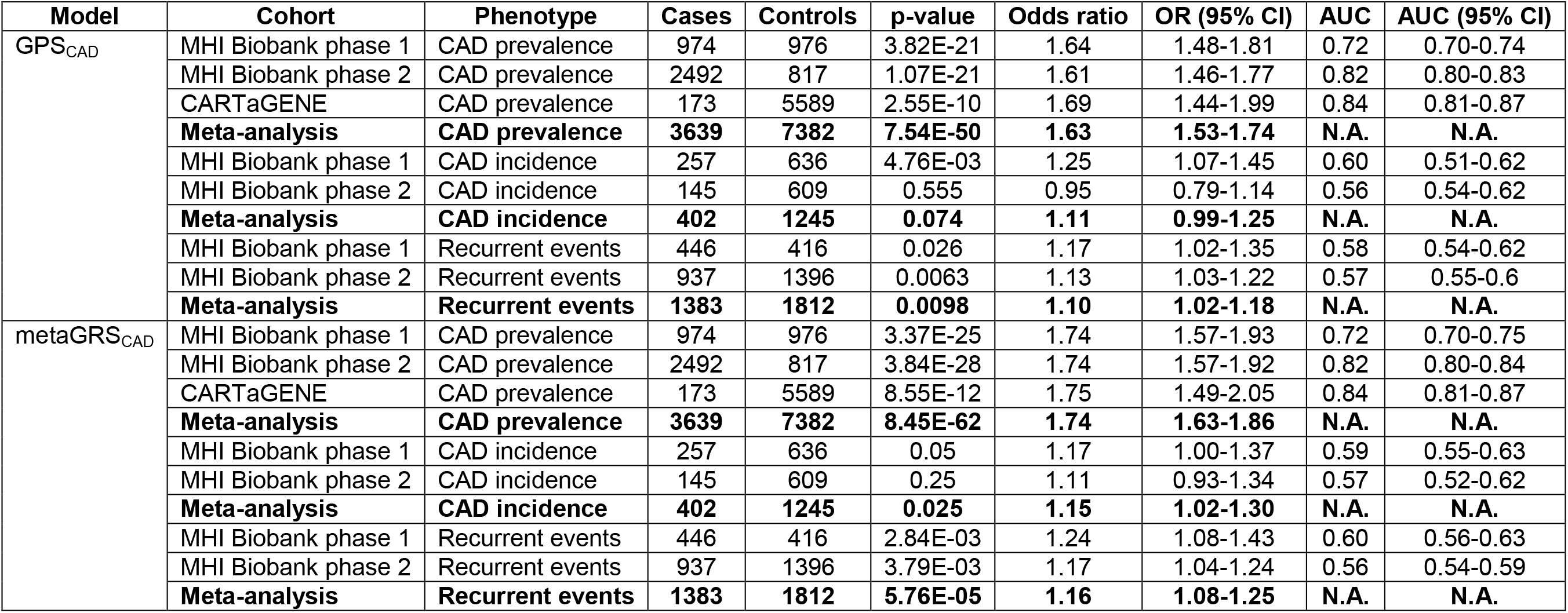
Association with and prediction of coronary artery disease by polygenic risk scores in three cohorts. N.A. = not applicable.

**Figure 1.**
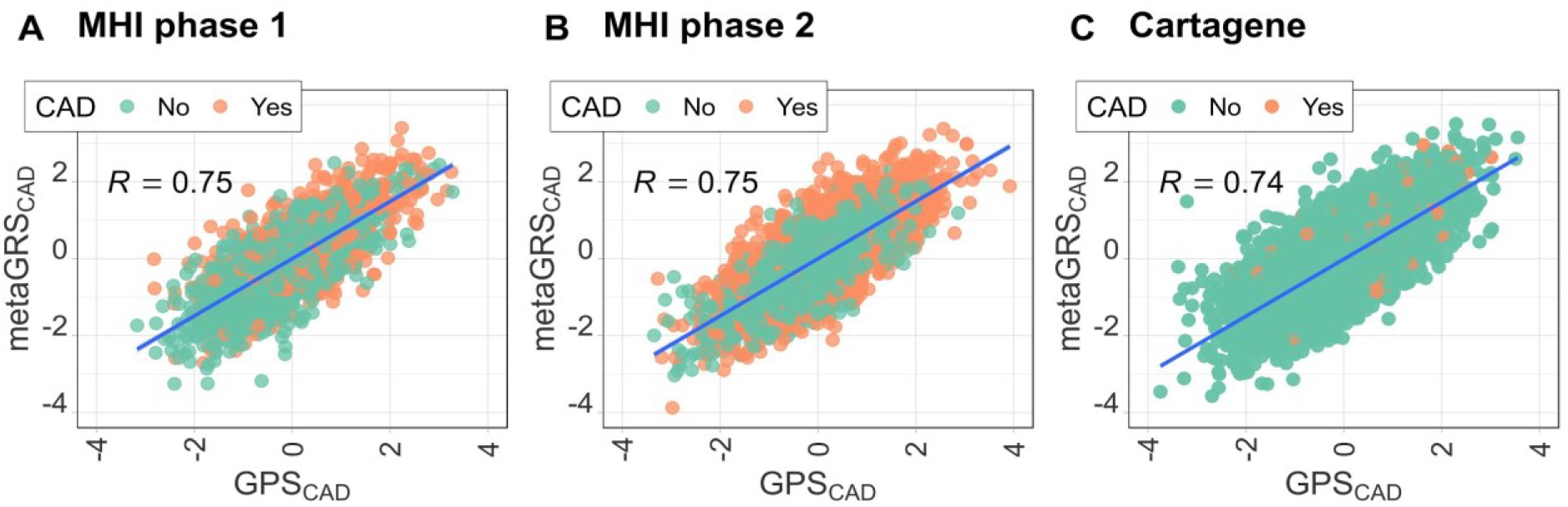
Correlation between normalized GPS_CAD_ and metaGRS_CAD_. The correlation between GPS_CAD_ and metaGRS_CAD_ in (**A**) the MHI Biobank phase 1 (Pearson’s *r* = 0.75, p-value <2×10^-16^), (**B**) the MHI Biobank phase 2 (Pearson’s *r* = 0.75, p-value <2×10^-16^), and (**C**) CARTaGENE (Pearson’s *r* = 0.74, p-value <2×10^-16^).

**Figure 2.**
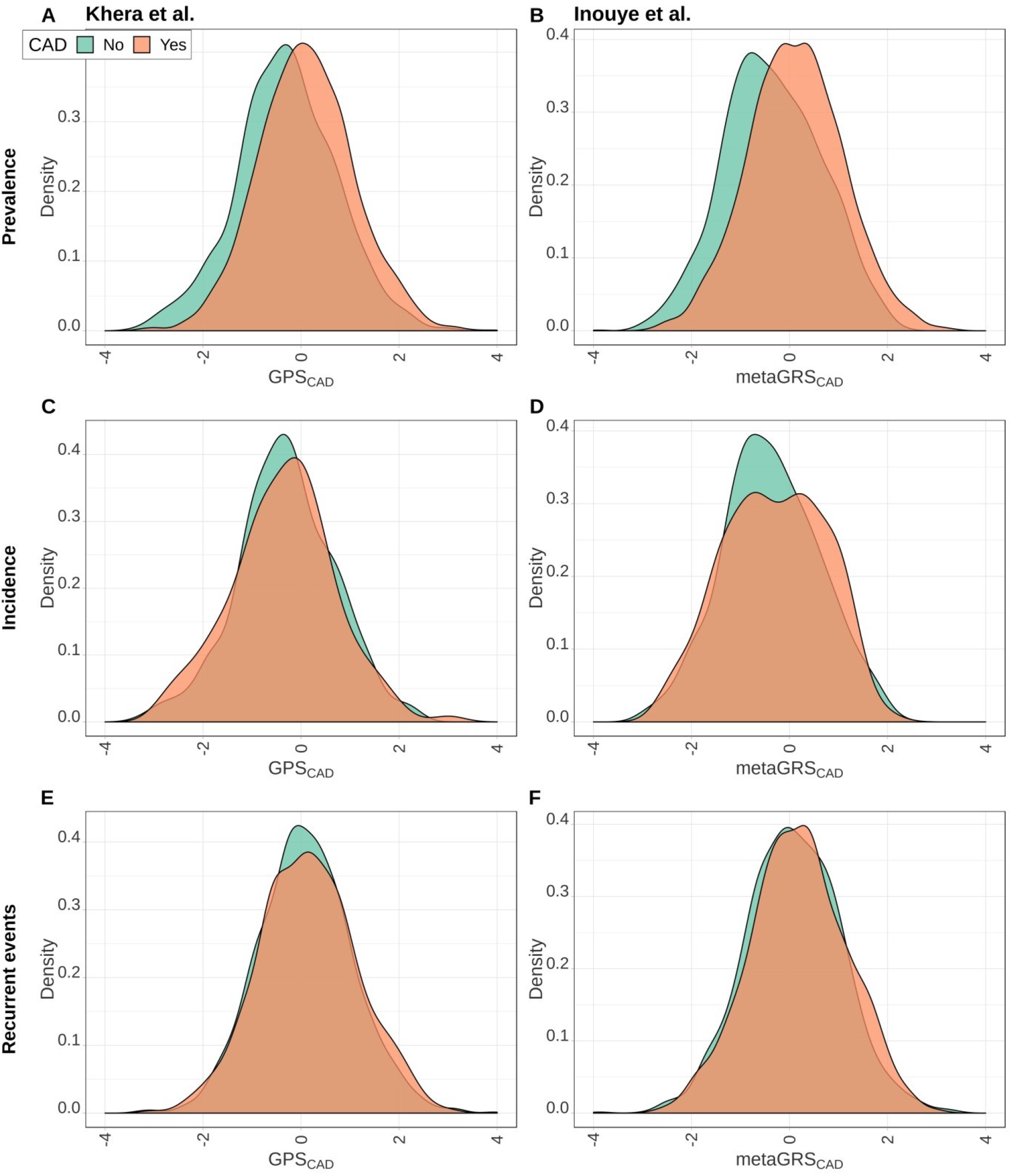
Distributions of GPS_CAD_ and metaGRS_CAD_ in the MHI Biobank phase 2 cohort. Distributions of the normalized polygenic risk score from Khera et al. (GPS_CAD_, left column) and Inouye et al. (metaGRS_CAD_, right column) in the MHI Biobank phase 2 data for prevalent (**A**,**B**), incident (**C**,**D**), and recurrent (**E**,**F**) coronary artery disease events.

### Estimation of CAD risk for *LDLR* delta > 15kb deletion carriers

Approximately 60% of FH cases in the French-Canadian population of Quebec are due to the delta > 15kb deletion of the *LDLR* gene ^22^. To compare the predictive power of CAD PRS with the impact of penetrant FH mutations on CAD risk in this population, we used whole-genome sequence (WGS) data available in 1,964 MHI Biobank participants to call copy-number variants at the *LDLR* locus ^19^. We identified a total of 14 heterozygous carriers of the *LDLR* delta > 15kb deletion (breakpoints: chr19:11,188,403-11,204,295 (hg19)). The estimated allele frequency in this cohort is 0.36%, which is in the range of the reported frequency (~0.03-0.38%) ^23,24^. In our dataset, the *LDLR* delta > 15kb deletion was associated with increased LDL-cholesterol levels (LDL-cholesterol: 1.34 mmol/L increase per copy of the *LDLR* deletion, p-value=1.2×10^-8^). When combining baseline and follow-up data, we found that 12 out of the 14 *LDLR* deletion carriers were CAD cases (odds ratio (OR)=3.30 and 95% confidence interval=0.72-15.2; p-value=0.13). Although this result is not statistically significant owing to our limited sample size, it allows us to estimate that French Canadians who carry a strong FH mutation are ~3 times more at risk to develop CAD. This provides a direct opportunity to identify the proportion of individuals at similar or increased risk for CAD based on their PRS. Using the distributions of GPS_CAD_ and metaGRS_CAD_, we estimate that 6-7% of the French-Canadian population is at the same or higher risk for CAD than carriers of the FH *LDLR* delta > 15kb deletion. This result is consistent with the estimate by Khera et *al*. that 8% of European-ancestry individuals in the UK Biobank have a PRS that confers comparable or higher CAD risk than rare FH mutations ^11^

### Prediction of incident and recurrent CAD

The MHI Biobank is a prospective hospital-based cohort with available regular follow-up clinical information collected. We took advantage of this design to also test the CAD PRS against incident and recurrent CAD events. Because genetic variants are present at birth, it can be argued that PRS analyses of late-onset diseases such as CAD are always prospective. However, analyses of clinical information collected retrospectively is subject to selection biases and thereby analysis of such information might impact the accuracy of the PRS. Inouye et *al*. had shown that metaGRS_CAD_ can identify incident cases in the UK Biobank ^12^. Among the 1245 controls at baseline with follow-up available in the combined MHI Biobank cohorts, 402 had a first CAD event between recruitment and follow-up (median follow-up time = 4 years (range = 5 weeks - 7.2 years)). GPS_CAD_ was not associated with incident CAD (OR=1.11, p-value=0.074), whereas the association between metaGRS_CAD_ and incident CAD was only modest (OR=1.15, p-value=0.025) (Table 2). The prediction of incident CAD by GPS_CAD_ and metaGRS_CAD_ was also markedly lower than for prevalent CAD (AUC=0.56-0.60) (Table 2). Of the 1812 CAD cases at baseline with follow-up information available, 1382 had a recurrent CAD event during the follow-up period (median follow-up time in years = 3.9 years (range = 1.1 - 7)). We found that GPS_CAD_ and metaGRS_CAD_, two PRS developed to predict primary CAD events, were also associated with recurrent CAD events (GPS_CAD_: OR=1.10, p-value=0.0098; metaGRS_CAD_: OR=1.16, p-value=5.76×10^-5^), although the AUC were relatively small (0.56-0.60) (Table 2).

## Discussion

Because PRS are simple and relatively inexpensive, their implementation in the clinical setting holds great promises. For CAD in particular, early detection could lead to simple yet extremely efficacious therapeutic interventions (*e.g.* statins, aspirin). Given this exciting possibility, we tested two recently developed CAD PRS in French Canadians recruited from population- and hospital-based cohorts. We validated previous findings that both GPS_CAD_ and metaGRS_CAD_ perform well for prevalent CAD cases. However, their performance was lower for incident and recurrent CAD in the MHI Biobank. Using the French-Canadian founder FH *LDLR* delta > 15kb mutation to calibrate CAD risk, we confirmed that PRS can identify about 6-7% of the population that is at equal or higher CAD risk than carriers of a FH monogenic mutation.

Our study raises a few interesting questions. Although it is appreciated that PRS do not transfer well between ancestral populations^25,26^, little is known about the transferability of PRS across populations within the same ancestry. Our results indicate that CAD PRS developed in European-ancestry individuals perform quite well in the genetically and environmentally homogenous French-Canadian population. How well these same PRS would predict CAD in a more diverse European-ancestry population, or in a population living in a very different environment, remain critical open questions for further investigation. Another important result from our analyses is the lower accuracy that these PRS have to predict incident or recurrent CAD cases when compared to prevalent CAD cases, highlighting the importance of the method used to create the PRS. GPS_CAD_ and metaGRS_CAD_ were built using mainly GWAS for prevalent CAD, and are therefore particularly suitable to predict prevalent CAD as opposed to incident or recurrent events. In particular, our analyses of incident and recurrent CAD were based on the MHI Biobank, which is a hospital-based cohort. Thus, it is possible that confounders such as the presence of co-morbidities and medications (*e.g.* anti-thrombotic) would impact PRS performance. It is important to clarify these differences in order to determine what factors in the study design and what ascertainment biases influence the PRS. Furthermore, an extension of our results implicates that GWAS that aim to specifically identify the genetic architecture of incident or recurrent CAD events might yield improved predictive power to calibrate risk score models over PRS based on CAD prevalence alone.

In conclusion, while it may still take some time before PRS become widely applicable in the clinic to predict CAD, their utility is likely to increase as the community continues to improve methods and gain access to large GWAS carried out in populations of different ethnic backgrounds. But true improvement in CAD prediction based on PRS will only occur if the scientific progress is mirrored by an effort to explain the strengths and limitations of this new biomarker to the medical community and the general population.

## Methods

### Data sources and availability

All participants have provided written informed consent and the project was approved by the ethics committee of the MHI. The low-pass whole-genome sequencing (WGS) dataset (phase1) and the genotyping dataset (phase 2) of the MHI Biobank have been previously described.^19,20^. Case status for CAD prevalence was defined as having a myocardial infarction (MI) or coronary artery interventions (coronary artery bypass grafting (CABG) or percutaneous coronary intervention (PCI)) before the first visit. Controls were selected to be free of MI, PCI, CABG, transient ischemic attack or stroke, peripheral vascular disease, congestive heart failure, and angina. We used the same clinical definitions to identify incident and recurrent CAD events in the MHI Biobank: participants who had a first-ever CAD event between baseline and follow-up were considered incident cases and participants who had at least a second CAD event during the same period were considered recurrent CAD cases. Average follow-up time was 4.2 years for phase 1 and 3.6 years for phase 2. We used GenomeStrip (v.2.0) with default parameters on the MHI Biobank phase 1 WGS data to identify participants who carry the known French-Canadian founder *LDLR* deletion ^27^. We excluded the 14 individuals who carry the *LDLR* delta > 15 kb deletion from all subsequent PRS analyses.

CARTaGENE (www.cartagene.qc.ca) is a population-based cohort of Quebec that includes individuals aged between 40 and 69 years ^21^. A subset of this cohort totaling 5762 samples were genotyped on the Illumina Infinium Global Screening Array (GSA). We used PLINK (version 1.9, https://www.cog-genomics.org/plink/1.9/) to apply the following quality-control filters: we excluded samples and variants with >5% missingness, variants out of Hardy-Weinberg Equilibrium (p-value<1×10^-6^), A/T and G/C variants, and variants with a minor allele frequency (MAF) <1%. Following these quality-control steps, we phased genotypes with ShapeIT v2.r790 and imputed missing genotypes on the Michigan Imputation Server (version 1.30.4) using the Haplotype Reference Consortium panel (Version r1.1 2016).^28,29^. Case-control status for CAD in CARTaGENE was defined using the same criteria than for the MHI Biobank (see above). Principal components for all three cohorts were calculated in PLINK using the *pca* function.

### Polygenic risk scores

The models for the two PRS (GPS_CAD_ and metaGRS_CAD_) used in this study are available online (see Web links below) ^11,12^. Genetic risk scores for all models were generated with PLINK (version 1.9) and the --score function to calculate the sum of the product of the dosage of effect alleles per variant weighted by the CAD effect size ^30^. We excluded variants with low imputation quality score (rsq <0.3). PRS were Z-score normalized and centered (mean=0, standard deviation=1) per dataset to facilitate interpretation of odds ratios. Detailed numbers for missing variants from all models can be found in **Table S1**.

### Statistical analysis

We performed all statistical analysis in R (version 3.5.0) ^31^. In the MHI Biobank datasets, we tested by logistic regression the association between CAD case-control status and PRS Z-scores correcting for age, sex, and the first four principal components. In the MHI Biobank phase 1 data, we also corrected for statins use (all MHI Biobank cases from phase 2 were on statin treatment). For the CARTaGENE data, we used a similar logistic regression model, correcting for age, sex, the first four principal components, and recruitment center. We calculated odds ratios per standard deviation of the GPS_CAD_ or metaGRS_CAD_ PRS, and considered p-value < 0.05 as significant. We calculated AUC using the pROC package in R ^32^. Meta-analysis was performed with the R metafor package using the coefficients and standard errors (SE) from the individual regression models with a fixed effect model fitted with the “FE” method ^33^. We tested the association between LDL-cholesterol levels and the *LDLR* deletion in RVTest using the *score test* function ^34^. For these analyses, we increased the LDL-C levels of dyslipidemic participants by 30% to account for the effect of statins.

## Funding information

This work was funded by the Canadian Institutes of Health Research (MOP #136979), the Heart and Stroke Foundation of Canada (Grant #G-18-0021604), the Canada Research Chair Program, Genome Quebec and Genome Canada, and the Montreal Heart Institute Foundation.

## Competing interests

The authors declare no competing interests.

## Acknowledgements

We thank all participants and staff of the André and France Desmarais MHI Biobank. Sequencing of the MHI Biobank samples (phase 1) was performed at the McGill University and Génome Québec Innovation Centre. Genotyping of the MHI Biobank samples (phase 2) was performed at the Université de Montréal Beaulieu-Saucier Pharmacogenomics Centre at the MHI. The authors would also like to thank the CARTaGENE staff for their support in validating phenotypes. We also thank Aikaterini Kritikou and Rafik Tadros for comments on an earlier version of this manuscript.

## Web links

Genetic risk score model for GPS_CAD_ Khera et al. Nature Genetics 2018: http://www.broadcvdi.org/informational/data

Genetic risk score model for metaGRS_CAD_ by Inouye et al. JACC 2018: https://figshare.com/articles/Coronary_Artery_Disease_CAD_MetaGRS/5748096

