## Supplementary_information for "Validation of genome-wide polygenic risk scores for coronary artery disease in French Canadians"

**Table S1.** Variants available from each cohort to calculate the two coronary artery disease polygenic risk scores. PRS = polygenic risk score used, n variants model = Total variants included in the model, variants scored = Percentage of variants from model that were present in data and used for scoring, variants missing = percentage of variants missing from scoring, rsq < 0.3 = number of variants that were excluded due to low imputation quality, other ALT = number of variants excluded due to different alternative allele than in reference model, missing = number of PRS model variants absent in dataset. N.A. = not applicable.

| Cohort | PRS | n variants model | variants scored (%) | variants missing (%) | rsq < 0.3 | other ALT | missing |
|---|---|---|---|---|---|---|---|
| MHI Biobank phase1 | $GPS_{CAD}$ | 6630150 | 94.01 | 5.99 | N.A. | 5394 | 391674 |
| MHI Biobank phase2 | $GPS_{CAD}$ | 6630150 | 93.04 | 6.96 | 47323 | 50 | 414406 |
| CARTaGENE | $GPS_{CAD}$ | 6630150 | 94.46 | 5.54 | 1475 | 3203 | 363401 |
| MHI Biobank phase1 | $metaGRS_{CAD}$ | 1745179 | 97.55 | 2.45 | NA | 616 | 42124 |
| MHI Biobank phase2 | $metaGRS_{CAD}$ | 1745179 | 96.16 | 3.84 | 33351 | 85 | 33595 |
| CARTaGENE | $metaGRS_{CAD}$ | 1745179 | 99.91 | 0.09 | 1475 | 50 | 0 |
